# Emergence of multiple spontaneous coherent subnetworks from a single configuration of human connectome-coupled oscillators model

**DOI:** 10.1101/2024.01.09.574845

**Authors:** Felipe A. Torres, Mónica Otero, Caroline Lea-Carnall, Joana Cabral, Alejandro Weinstein, Wael El-Deredy

**Affiliations:** Biomedical Engineering, Universidad de Valparaíso, Valparaíso, Chile; Brain Dynamics Laboratory, Universidad de Valparaíso, Valparaíso, Chile; Facultad de Ingeniería, Arquitectura y Diseño, Universidad San Sebastián, Santiago, Chile; Centro Científico y Tecnológico de Excelencia Ciencia & Vida, Universidad San Sebastián, Santiago, Chile; School of Health Sciences, Faculty of Biology, Medicine and Health, Manchester Academic Health Science Centre, University of Manchester, Manchester, UK; Life and Health Sciences Research Institute, Minho University, Minho, Portugal; ValgrAI, Valencian Graduate School Research Network of Artificial Intelligence, Spain; Department of Electronic Engineering, School of Engineering, Universitat de València, Spain; Advanced Center of Electrical and Electronics Engineering, Valparaíso, Chile

## Abstract

Large-scale brain models with biophysical or biophysically inspired parameters generate brain-like dynamics with multi-state metastability. Multi-state metastability reflects the capacity of the brain to transition between different network configurations and cognitive states in response to changing environments or tasks, thus relating to cognitive flexibility. To study this phenomenon, we used a Kuramoto network of oscillators corresponding to a human brain atlas of 90 nodes, each with an intrinsic frequency of 40 Hz. The network’s nodes were interconnected based on the structural connectivity strengths and delays found in the human brain. We identified global coupling and delay scale parameters corresponding to maximum spectral entropy, a proxy for maximal multi-state metastability. At this point, we show that multiple coherent (functional) sub-networks spontaneously emerge across multiple oscillatory modes, and persist in time for periods between 140 and 4389 ms. Most nodes in the network exhibit broad frequency spectra away from their intrinsic frequency, and switch between modes, in a manner similar to that reported in empirical resting-state neuroimaging data. We suggest that the obtained dynamics at maximum metastability is a suitable model of the awake brain. Further, we show that global coupling and delay scale parameters away from maximum metastability yield dynamical features similar to other brain states such as sleep and anesthesia. Therefore, spectral entropy also correlates with wakefulness and the synchrony of functional networks.

## Introduction

Large-scale brain network models based on biologically plausible local dynamics and interconnected using structural connectivity data derived from human imaging studies have offered a principled framework for studying the emergence of global brain dynamics. Simulated brain activity based on these models resembles realistic neural processes and can be used to make predictions about neurological disorders (***Cabral et al., 2011***; ***Breakspear, 2017***; ***Menon, 2011***).

Insights from large-scale brain network models include efficient information processing due to small-world network properties (dense local and sparse long-range connections) (***Black and Rogers, 2020***; ***Bullmore and Sporns, 2009***); the presence of hubs for information integration and transfer (Bassett and Sporns, 2017), “rich clubs” of highly interconnected hub regions for rapid communication (Bullmore and Sporns, 2012), and networks associated with specific cognitive functions, such as the default mode network for introspection and the salience network for attention and emotion (***Lydon-Staley et al., 2019***).

One of the most revealing insights from large-scale brain network models is the dynamic nature of the brain’s functional networks and their rapid reconfiguration, spontaneously or in response to tasks and stimuli (***Finc et al., 2020***), and that this dynamic functional connectivity (FC) is essential for adaptability and cognitive flexibility (***Capouskova et al., 2023***). This capacity for fast transitioning between different functional network configurations that persist over time in response to cognitive and environmental demands is called metastability (Tognoli and Kelso, 2014).

Inter-regional coupling strength and connection delays play important roles in brain metastability. The strength of coupling between different brain regions affects the degree of synchronization between them. Stronger coupling promotes synchronization, i.e., regions become more coordinated and aligned for longer durations, while weaker coupling results in desynchronization or more independent neural activity.

The balance between strong and weak coupling is essential for achieving both integration (communication between brain regions) and segregation (specialized processing within regions) (***Bian et al., 2021***). Metastability involves maintaining this balance, allowing the brain to transition between integrated and segregated states as needed for different cognitive functions (***Orio et al., 2018***).

On the other hand, connection delays, or the time it takes for signals to propagate between brain regions, introduce temporal dynamics into the brain’s patterns of FC and contribute to the emergence of metastable states as the brain navigates through different network configurations (***Nakagawa et al., 2014***; ***Petkoski and Jirsa, 2019***; ***Williams et al., 2023***). The timing of transitions between states can depend on the combination of delays within the network (***Cabral et al., 2022***).

The interplay between coupling strength and connection delays is complex and can result in various dynamic states. When brain regions are weakly-coupled with long delays, their activity is independent. As the coupling strength increases and the delay reduces, the system transitions towards fully synchronized dynamics.

The dynamic interplay between these factors allows the brain to balance synchronization and desynchronization, integration and segregation, the temporal coordination of neural activity, and the peak synchronization frequency (***Cabral et al., 2022***). Understanding how coupling strength and connection delays influence brain metastability is a crucial aspect of neuroscience (***Abeysuriya et al., 2018***; ***Deco et al., 2014***; ***Wildie and Shanahan, 2012***).

The emergence of functional coherent subnetworks at different fundamental frequencies during the resting-state regime of the brain occurs spontaneously. A similar emergence of subnetworks or oscillatory modes is also expected from a multi-state metastability regime. It is important to highlight that the coherent subnetworks in the brain also present a spatial preference for a subset of the brain regions and a preferred fundamental frequency (***Cabral et al., 2022***; ***Vidaurre et al., 2018***).

This paper aims to demonstrate the emergence of multiple oscillatory modes, their corresponding coherent functional networks, and their persistence in time from a network of Kuramoto oscillators. The network nodes correspond to brain regions connected according to the Automatic Anatomical Labelling (AAL90) Atlas (***Tzourio-Mazoyer et al., 2002***) using a single configuration of coupling and delay parameters.

We assume spectral entropy to be a proxy for metastability and identify the set of coupling and delay parameters that maximises this metric. We also present the emergent spectrum and metastable dynamics of the brain regions at that set of parameters. Further, we characterize the FC at specific frequency bands, showing the emergence of tempo-spatial and frequency-specific functional subnetworks. Finally, we compute subnetworks’ temporal features searching for differences in their fractional occupancy and duration distributions.

We show a correlation between spectral entropy and wakeful brain states. When spectral entropy reaches its peak, the resulting coherent subnetworks from our model resemble the expected networks observed in empirical human resting-state EEG. This resemblance encompasses both spatial and temporal features.

## Results

The emerging dynamics from delayed interactions among identical oscillators exhibit a spectrum with frequency components different from the single intrinsic frequency of each individual oscillator. We are interested in how many different peaks and how much broader this emergent spectrum becomes when we vary the global parameters: the global coupling coefficient *K* and the mean conduction delay MD.

### Parameters set for metastable dynamics

Multi-state metastable dynamics have at least two attractor basins by definition. This implies that the system’s average spectral profile will exhibit a shape distinct from a single frequency peak (see ***Figure 1***-figure Supplement 1, for examples of the average spectral profiles resultant from the Kuramoto model). The relevant frequency components increase with the number of attractor basins in the system.

**Figure 1.**
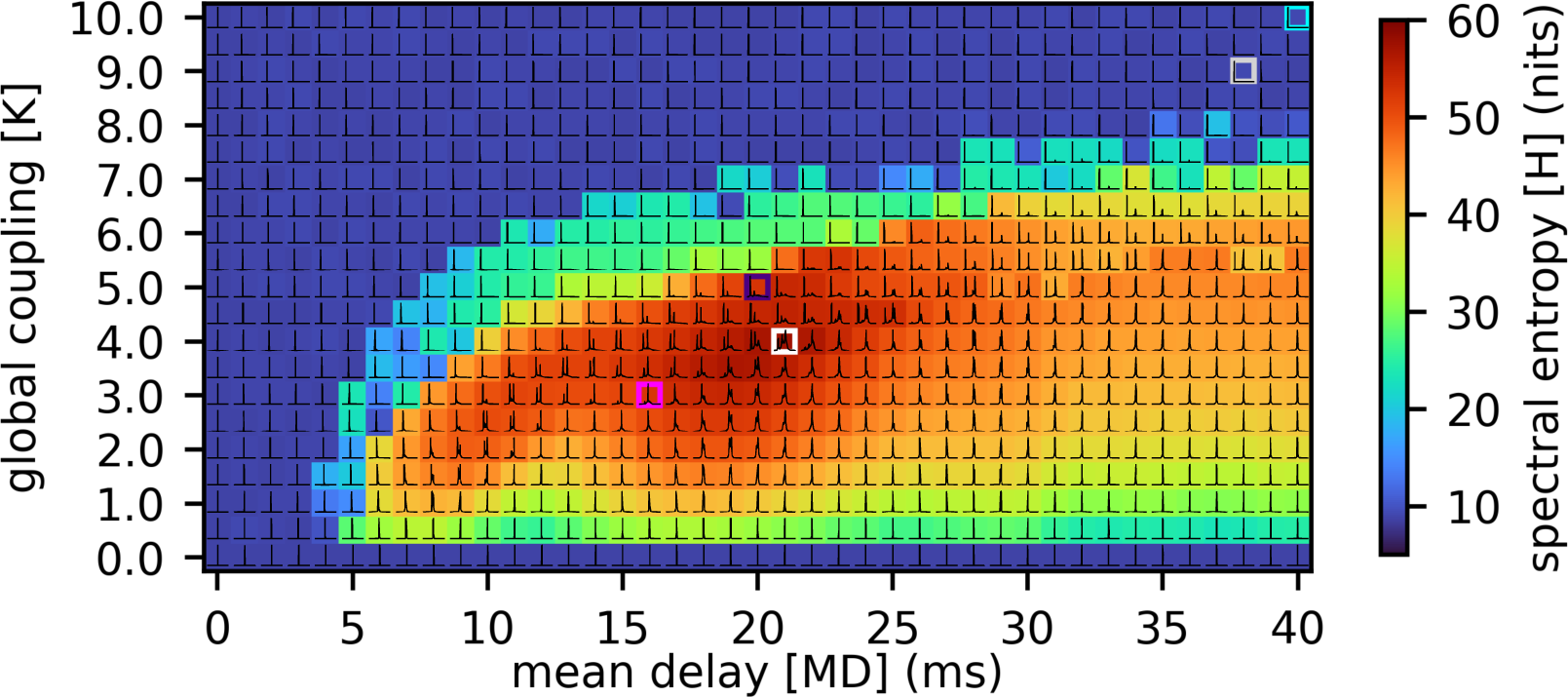
Spectral entropy as a metric of metastability from the parameters space exploration of global coupling, *K*, and mean delay MD. The white square marks the maximum value obtained for the spectral entropy at *K* = 4 and MD = 21 ms. The dark-blue square marks the best correlation fit with resting-state EEG spectrum at *K* = 5 and MD = 20 ms, and the magenta square marks the best fit from *Cabral et al*. (2014) at *K* = 3 and MD = 16 ms. The cyan square marks the best fit with anesthesia EEG at *K* = 10 and MD = 40 ms. The grey square marks the best fit with slow-wave sleep EEG at *K* = 9 and MD = 38 ms. Figure 1—figure supplement 1. Normalized average spectrum for three values of *K* = {2, 4, 6} and for different values of mean delay.

There is no specific numerical quantity that directly measures metastability. Instead, various quantitative and at times qualitative metrics are employed to assess the features of brain function associated with metastability (***Shanahan, 2010***). Here, we use the spectral entropy as a proxy for metastability while varying the global coupling *K* and the mean conduction delay MD. ***Figure 1*** shows a colored heatmap of the corresponding spectral entropy for each set of parameters in our model. ***Figure 1*** also includes the average spectrum for each set of parameters.

The maximum value obtained for the spectral entropy is highlighted with a white square corresponding to *K* = 4 and MD = 21 ms. This set of parameters also presents low synchrony assessed by the average of the Kuramoto order parameter (KOP) (***Figure 1***-figure Supplement 2 A) and it is not the maximal set of parameters for the KOP standard deviation (***Figure 1***-figure Supplement 2 B). The best fit with FC profiles of resting-state MEG is marked with a magenta square (*K* = 3, MD = 16 ms) as found in ***Cabral et al. (2014)***. The point of the best fit correlating the Kuramoto model average spectrum to resting-state EEG average spectrum is marked with a dark-blue square (*K* = 5, MD = 20 ms) (***Rivera et al., 2023***). The point of best correlation with anesthesia EEG spectrum is marked with a cyan square (*K* = 10, MD = 40 ms) (***Liu et al., 2018***), and the correlation with slow-wave sleep EEG spectrum is marked with a grey square (*K* = 9, MD = 38 ms) (Devuyst, 2005). The spectra of resting-state, anesthesia, and slow-wave sleep EEG are corrected by 1/f to remove the aperiodic component before correlating with the model spectra.

The coupling value and mean delay in anesthesia and slow-wave sleep are higher than in the resting-state case. Thus, these brain stages have higher synchrony but lower spectral entropy.

**Figure 1—figure supplement 2. A.** Average Kuramoto order parameter, the synchrony of the entire system. B. The standard deviation of the Kuramoto order parameter as another metastability metric. C. Peak frequency of the average spectrum.

### Emerging spectrum and time switching between attractor basins

The average spectrum of the system indicates the frequencies in which the system’s power is highly concentrated. We assume that the frequency peaks are associated with the existence of limit-cycle attractors. We focus on the system activity at the frequency peaks indicated in Figure 2 A because these correspond to the major EEG and MEG wavebands found in human brain data (***Herrmann et al., 2016***; ***Mellem et al., 2017***). We band-pass filtered the time-series data using a bandwidth of 1 Hz around these peak frequencies. Then, we extracted the envelopes to determine the dominance of every peak frequency at each time point (see Methods and Materials-Selection of envelopes threshold for details of fractional occupancy calculation by thresholding the envelopes).

**Figure 2.**
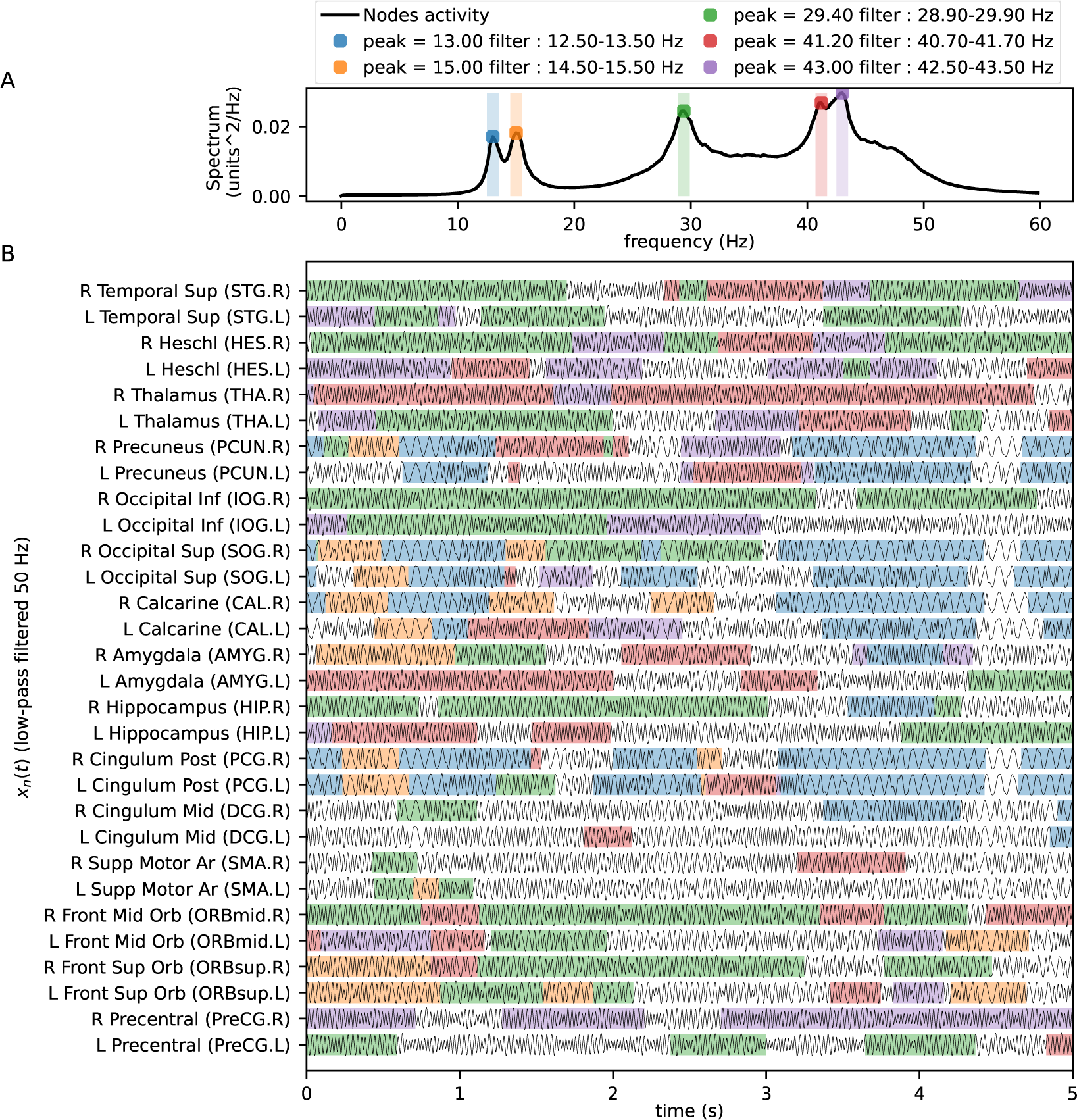
A. The average spectrum of the ninety nodes with *K* = 4 and MD = 21 ms. B. Time-series of the nodes’ activity filtered below 50 Hz. The shaded regions highlight when the nodes’ envelopes pass a threshold = 0.232 for each frequency band. As the maximum possible envelope amplitude is 1, the selected threshold (for threshold selection details, see Methods and Materials-Selection of envelopes threshold) indicates when at least 5.3% of the node’s instantaneous power is present at each narrow frequency band. Figure 2—figure supplement 1. Fractional occupancy of the ninety nodes at each frequency band. Figure 2—figure supplement 2. Average spectrum and time-series of the nodes using *K*=8, MD=37 ms.

As the spectrum corresponds to the set of parameters that determine maximum metastability, the visualization of the time excerpts of the Kuramoto model simulations in Figure 2 B shows how some nodes ’switch’ between attractors (see, for example, simulations from nodes relating to the brain areas Right Precuneus and Right Heschl). These nodes have low variance in their fractional occupancy at each peak frequency (see Figure 2-figure Supplement 1). On the other hand, some nodes have a preferred frequency most of the time (see, for example, Right Occipital Inferior). We note here that some nodes have most of their power in frequencies other than the frequency peaks defined here, i.e., the power of their oscillations in these bands did not pass the occupancy threshold (see, for example, Left Cingulum Mid).

On the other hand, the Kuramoto dynamics from a set of parameters that determine lower spectral entropy does not present several peak frequencies (see Figure 2-figure Supplement 2 A), neither “switch” between attractors (see Figure 2-figure Supplement 2 B).

### Functional sub-networks emerging at each frequency peak

FC relates to pairs of brain regions with correlated activities during a determined period (***Friston, 1994***). We use sliding windows with an overlap of 50% to determine the FC matrix by examining the spectral coherence at each frequency peak. Figure 3 shows the subnetworks with higher fractional occupancy at each peak frequency (see Methods and Materials-FC matrices clustering for clustering details, see Figure 3-figure Supplement 1 for region labels and positions reference). We found that emerging subnetworks at lower frequencies (Figure 3 A-B) have a greater number of interhemispheric connections than the higher frequency subnetworks (Figure 3 C-E).

**Figure 3.**
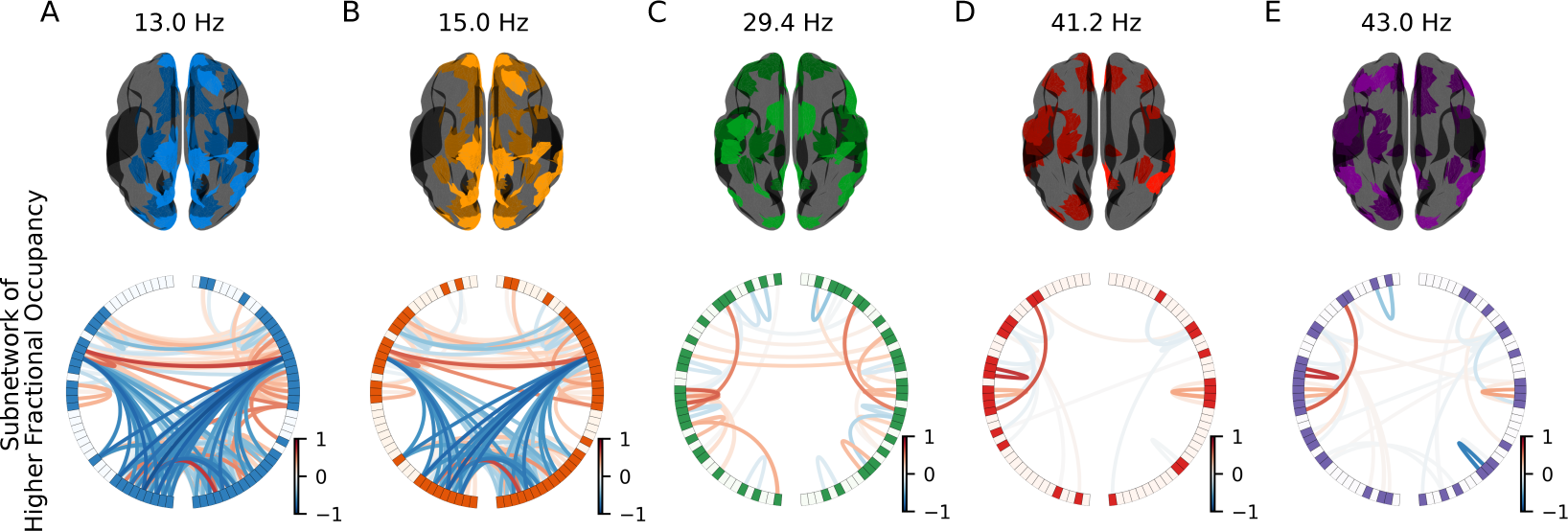
Functional subnetworks with the maximal fractional occupancy for each frequency peak (A-E). The average spectral coherence FC matrix of each cluster is masked by the structural connectivity matrix, and only the highest 10% of the connections are shown. The color of the brain regions corresponds to the colors previously used for each frequency band. The color of the edges in the circular network plot indicates the imaginary part of the coherence (*icoh*). *Figure 3*-*figure Supplement 2* shows the real part of the coherence. Figure 3—figure supplement 1. Structural connectivity and region degrees. The region labels correspond to the region in the same position of *Figure 3* and *Figure 3*-*figure Supplement 2*. The plotted edges are only the 10% higher connection weights. The gray color level of each region corresponds to the region degree. Figure 3—figure supplement 2. Real part of the edges coherence. The edges of the functional subnetworks have lower coherence as the frequency increases. Figure 3—figure supplement 3. Regions participating in all the obtained subnetworks. Using the original node order of the AAL90 matrix. Figure 3—figure supplement 4. Regions participating in all the obtained subnetworks, sorted from the least to the most functional connected node. Figure 3—figure supplement 5. Clustering score. The two clustering criteria are detailed in Methods and Materials section. The number of clusters for each frequency peak is the *M* value with the highest inter-trial correlation of the clusters centroids or where an elbow occurs. We also keep a high inter-trial correlation of the fractional occupancy. We selected 4, 5, 4, 7, and 7 subnetworks for the 13, 15, 29.4, 41.2, and 43 Hz frequency peaks, respectively.

The functional subnetworks at 13 Hz and 15 Hz (Figure 3 A-B) resemble the Default Mode Network (DMN) (***Buckner et al., 2008***; ***Vidaurre et al., 2018***). These include the medial prefrontal cortex (ORBsupmed, SFGmed, REC), the posterior cingulate cortex (PCG), the cuneus (CUN), the precuneus (PCUN), the angular gyrus (ANG), and the thalamus (THA). Note that the medial regions are connected to the occipital regions with a negative value of the imaginary coherence, while the interhemispheric connections have positive imaginary coherence. This is an example of anticorrelated, but coherent, simultaneous activity.

The subnetwork that emerges at 29.4 Hz (Figure 3 C) resembles a combination of the salience network with the sensory systems (***Fox et al., 2006***). This subnetwork includes the right insula (INS), the anterior cingulate cortex (ACG), the posterior cingulate cortex (PCG), the occipital superior gyrus (SOG - visual), the supplementary motor area (SMA), the postcentral gyrus (PoCG - motor), the superior temporal gyrus (TPOsup - auditory), and the middle temporal gyrus (TPOmid - visual).

The subnetworks at 41.2 and 43 Hz (Figure 3 D-E) resemble the fronto-parietal network (***Dosenbach et al., 2007***; ***Marek and Dosenbach, 2018***), which includes the frontal superior gyrus (SFG), the frontal inferior gyrus (IFG), the inferior parietal lobule (IPL), and the caudate (CAU), and the inferior temporal gyrus (ITG). However, it seems that the networks also include part of the salience network, given that they include the insula (INS) and the right posterior cingulate cortex (PCG).

The putamen (PUT), the pallidum (PAL), and the right caudate (CAU) require a mention, as they participate in all the subnetworks at all the five frequencies. On the other hand, the imaginary coherence of the right precentral gyrus (PRECg), left frontal inferior operculum (IFGoperc), and left supramarginal (SMG) do not participate in any functional network, as they do not have enough coherence with other regions at the five frequencies.

The additional subnetworks at each frequency band share a “core” of participant brain regions (Figure 3-figure Supplement 3 and Figure 3-figure Supplement 4). The regions “out of the core” are recruited/excluded at different times with as many different connectivity patterns such as the number of detected subnetworks.

### Fractional occupancy and duration of the frequency-related sub-networks

Using the sliding windows, we can calculate the fractional occupancy and duration of the subnetworks in each frequency band using the window length. The subnetwork’s fractional occupancy comes from the number of time windows belonging to each cluster in each frequency peak. The violin plots in Figure 4 A show the occupancy of the subnetworks for each frequency peak. There are statistical differences between certain groups at 13, 15, and 29.4 Hz (Kruskal-Wallis H-test p<0.05). All subnetworks at 13 and 15 Hz are pair-wise statistically different from each other (p<0.01, Wilcoxon t-test, Bonferroni corrected for multiple comparisons). Almost all subnetworks at 29.4 Hz also are pair-wise statistically different.

**Figure 4.**
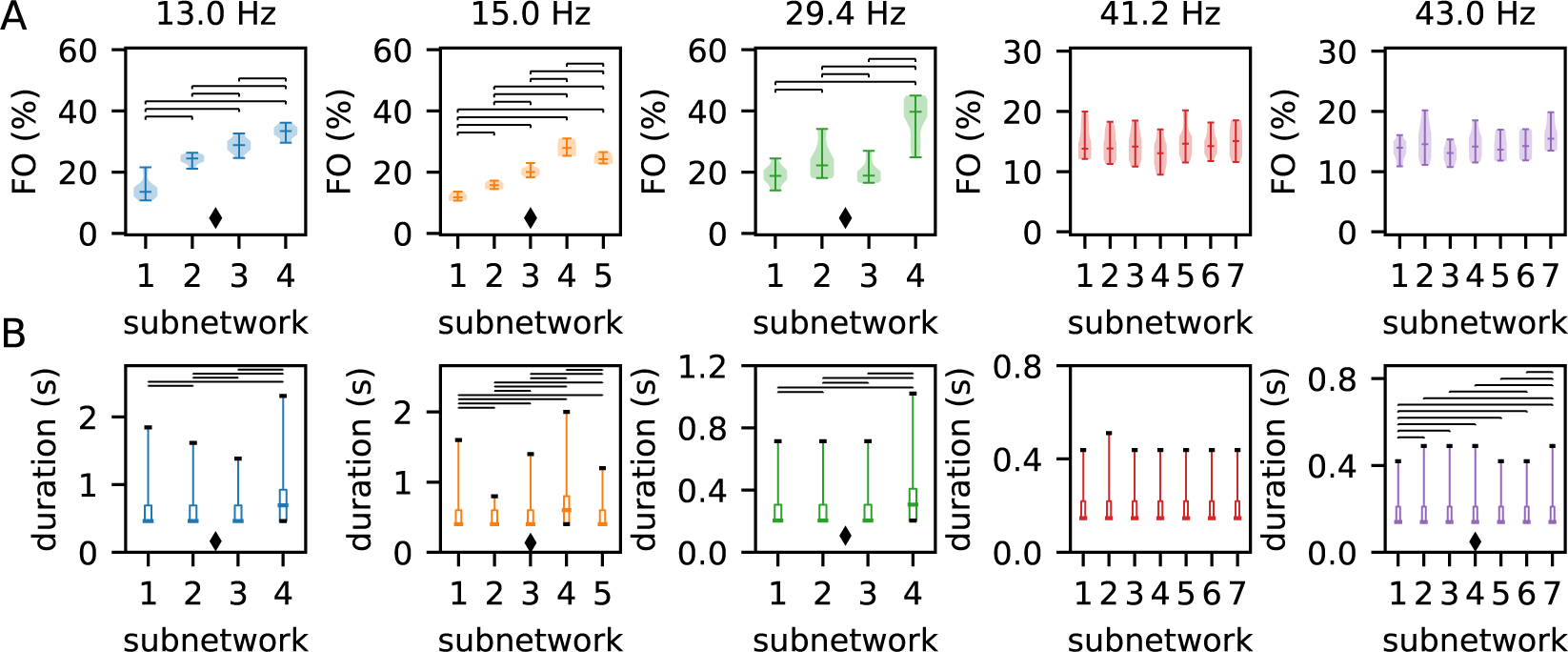
A. Fractional occupancy of the spectral coherence FC sub-networks. B. Duration distributions of the coherent sub-networks. Each vertical panel corresponds to the previously defined frequency bands. A diamond below the distributions indicates multi-group statistical difference (Kruskal-Wallis H-test p<0.05 in A, p<0.01 in B). The horizontal lines above the distributions indicate a statistically significant difference between the indicated pairs (p<0.01 Wilcoxon t-test). Figure 4—figure supplement 1. Excerpt of subnetworks occurrence. A. Occurrence of the subnetworks at each peak frequency along time. At each time point, there is one subnetwork of each peak frequency. B. Single time series of the overlapped subnetworks by relabeling the co-occurrences (There are a possible 3920 combinations from the 26 subnetworks at the five peak frequencies). C. Distribution of durations for overlapped subnetworks.

The median of the maximal fractional occupancy is 33.3% at 13 Hz and 27.8% at 15 Hz. The median of the maximal fractional occupancy at the medial frequency at 29.4 Hz is 39.7%. The median of the maximal fractional occupancy is 15.1% at 41.2 Hz (subnetwork 7), and 15.4% at 43 Hz (subnetwork 7).

Although the subnetworks’ occurrences are similar in their fractional occupancy at the two higher frequency peaks, it is possible to select one subnetwork with the highest fractional occupancy at each peak frequency.

The boxplots in Figure 4 B present the duration of the individual appearances of the functional subnetworks. Using six-cycle windows, we obtain a longer median duration for the lower frequency peaks. Interestingly, there are significant differences (p<0.01, Wilcoxon t-test, Bonferroni corrected for multiple comparisons) in the subnetworks duration between almost all the pairs of subnetworks at the three lower frequencies. At higher frequencies, there are fewer significant differences in the distribution of the subnetworks’ duration between all pairs. As multiple-group, there are statistical differences (Kruskal-Wallis H-test p<0.01) for four of the five peak frequencies. The exception in multiple-group comparison is at 41.2 Hz.

The maximal duration at each frequency are 4.389 s (subnetwork 4), 3.8 s (subnetwork 4), 2.856 s (subnetwork 4), 1.022 s (subnetwork 7), and 0.98 s (subnetworks 4 and 7), respectively.

## Discussion

### Model choice: Kuramoto network as a large-scale brain model

The Kuramoto model assigns a key role to the oscillators’ coupling. This model becomes a whole-brain model when the coupling corresponds with a human brain connectome inheriting the coupling strengths and connection delays (***Honey et al., 2007***, 2009; ***Ghosh et al., 2008***; ***Deco et al., 2009***). The Kuramoto model simplifies the dynamics of each region to a unitary amplitude harmonic oscillator. This simplification contrasts with biophysically-inspired oscillatory dynamics (***Abeysuriya et al., 2018***) or amplitude-varying oscillators (***Cabral et al., 2022***), but provides a mechanistic explanation of oscillatory phenomena in the human brain cortex (***Breakspear et al., 2010***; ***Cabral et al., 2011***, ***2014***; ***Daffertshofer and van Wijk, 2011***; ***Daffertshofer et al., 2018***). Not surprisingly, the Kuramoto model was proposed from the beginning by ***Kuramoto (1975)*** as a plausible model of the brain alpha-rhythm.

### Metric for multi-state mestastability

#### Emergence of metastable dynamics

Our results show the emergence of different sub-networks exhibiting a preferred frequency. Similar to the work of ***Roberts et al. (2019)*** and Cabral et al. (2022), the subnetworks are visited in sequence, and none of them persist indefinitely (Figure 2 B), suggesting that the simulated dynamics exhibit metastable behaviour. It is worth mentioning that we used conduction delays informed by the structural connectivity (***Cabral et al., 2014***, ***2022***), in contrast to the homogeneous delay used in ***Roberts et al. (2019)***. Also, we used a single natural frequency of 40 Hz for all nodes rather than retrieving different natural frequencies for each node from a distribution as in the work of ***Pang et al. (2021)***. Finally, we highlight that our model (***Cabral et al., 2014***) does not include any external stimulation or intrinsic noise.

#### Measuring the metastability

The standard deviation of the Kuramoto order parameter, which has been used as a proxy measurement of metastability (***Shanahan, 2010***; ***Wildie and Shanahan, 2012***), indicates the global variation between synchronized and non-synchronized states. However, a multi-state metastable system does not only interchange between synchronized/not-synchronized states within a single average frequency. Instead, it shows different synchronization levels around multiple peak frequencies. The spectral entropy of each node indicates the richness of its dynamics and can be thought of as a measure of the information contained in its frequency spectrum. The sum of the entropy spectrum from all the nodes gives a global measurement of the diversification of the states/modes of the system (***Vakkuri et al., 2004***; ***Rezek and Roberts, 1998***; Inouye et al., 1991).

Thus, we start by determining the maximum value for a specific combination of the global coupling and mean conduction delay.

We select the point in the model’s parameter space that maximizes the spectral entropy to ensure greater diversity in the dynamics of the simulated time series. However, using entropy to measure the “richness” of emerging dynamics in the brain or any other complex system could be ill-determined given that the entropy value is maximal for white noise (***Papo, 2016***). Nonetheless, it is an adequate metric for diverse emerging dynamics when all the nodes start with a single peak in their spectrum by imposing the natural frequency on the model, i.e., the minimum possible entropy. We suggest the maximum value of spectral entropy is limited by the structural features of the employed network, as expected from geometrical constraints on brain dynamics ***Pang et al. (2021)***.

### Emergence of coherent subnetworks

We argue that the different emerging frequency peaks correspond to several limit-cycle attractor basins. Following this presence of multiple frequencies, spectral coherence is an adequate measurement of frequency-specific synchronization (***Srinivasan et al., 2007***; Prado et al., 2023a). Moreover, spectral coherence indicates the presence of phase-locking pairs at all frequency ranges. We limited the analysis and results to the peak frequencies of the average spectrum (Figure 2 A), but this approach could be used in other central frequencies. However, some caution must be taken when using frequency-domain measurements. The main disadvantage of coherence with respect to the instantaneous Kuramoto order parameter is the need to use time windows. We use a different time window length for each frequency peak, and an overlap of 50% to counteract the temporal disadvantage and to get a higher temporal resolution. Then, the obtained subnetworks represent different phase-locking patterns at each peak frequency.

### Regions participate in different networks

We present the subnetworks with highest participation during the simulation period Figure 3 at the five peak frequencies. The coherence subnetworks at the two lower frequencies resemble the DMN. The subnetworks of the middle frequency (29.4 Hz) are a combination of the salience network and sensory networks. Finally, the subnetworks at the two higher frequencies resemble the fronto-parietal or central executive network. These networks are also called the three core neurocognitive networks, and dysregulation of these networks correlates with various neuropathologies (***Menon, 2011***).

The basal ganglia regions: pallidum (PAL) (structural features on left hemisphere: strength *I*_*n*_ = 88.27, degree *k*_*n*_ = 53; right: *I*_*n*_ = 82.44, *k*_*n*_ = 58. See Methods and Materials-Structural features of human-connectome, for details of *I*_*n*_ and *k*_*n*_ computation), putamen (PUT) (left: *I*_*n*_ = 130.84, *k*_*n*_ = 57; right: *I*_*n*_ = 141.29, *k*_*n*_ = 62), and the right caudate (CAU) (*I*_*n*_ = 97.46, *k*_*n*_ = 63) participate in all the subnetworks at all peak frequencies. But their participation is mainly via connections between themselves. However, the left caudate (*I*_*n*_ = 77.77, *k*_*n*_ = 57) does not participate in the networks of 29.4 Hz and 41.2 Hz. Perhaps the lack of the left caudate to complete the basal ganglia is a threshold artifact or could come from the lower structural connection strength than the other basal ganglia regions. In any case, the predominant presence of the basal ganglia indicates the importance of including sub-cortical regions to study brain dynamics (***Chakravarthy et al., 2010***). According to the presence of brain regions in each subnetwork (Figure 3-figure Supplement 4), the thalamus (THA) (left: *I*_*n*_ = 89.06, *k*_*n*_ = 64; right: *I*_*n*_ = 79.88, *k*_*n*_ = 64), right insula (INS) (*I*_*n*_ = 93.51, *k*_*n*_ = 43), right posterior cingulum gyrus (PCG) (*I*_*n*_ = 149.00, *k*_*n*_ = 61), and left anterior cingulum gyrus (ACG) (*I*_*n*_ = 124.84, *k*_*n*_ = 42) are present in almost all the subnetworks at all peak frequencies.

These regions present connections between them and other regions. Particularly, the cingulum regions present interhemispheric connections. The active presence of cingular regions in networks of resting-state dynamics has been demonstrated to be correlated with the anatomy of the cingulum bundle (***Skandalakis et al., 2020***; ***Fujiwara et al., 2007***; Van Den Heuvel et al., 2008).

This bundle, or collection of white matter tracts, connects regions of the frontal lobe with the precuneus (PCUN), posterior cingulate cortex (PCG), thalamus (THA), hippocampus (HIP), and parahippocampus (PHG) (***Bubb et al., 2018***).

The remaining 80 regions are absent in the subnetworks for at least one peak frequency. These regions are recruited by the subnetworks at different peak frequencies with preference in the effective frequencies of the region, which result from the interaction of all the ninety nodes.

There are still open questions at the functional level: Could the activity of the functional hub (basal ganglia, thalamus, insula, and cingulum gyrus) modify the recruitment of the other 80 regions? What is the result of malfunction/disconnection of one node of the functional hub?

Regarding the relation with structural features of the regions, note that the right PCG has the highest strength *I*_*n*_ = 149.00 but this region does not participate in all the subnetworks. Neither, the thalamus regions in both hemispheres having the highest degree *k*_*n*_ = 64 participate in all the subnetworks.

On the other hand, the right precentral (PreCG) (*I*_*n*_ = 65.94, *k*_*n*_ = 26), the left frontal inferior operculum (IFGoperc) (*I*_*n*_ = 82.88, *k*_*n*_ = 28) and the left supramarginal (SMG)(*I*_*n*_ = 59.10, *k*_*n*_ = 22) are absent from all the subnetworks. However, the region with lowest strength *I*_*n*_ = 46.78 is the right paracentral lobule (PCL) participating in five subnetworks. The region with lowest degree *k*_*n*_ = 18 is the right rolandic operculum (ROL) participating also in five subnetworks. Thus, we can expect more participation in functional subnetworks from the highly connected regions, but we can not predict how much greater it may be.

### Temporal features of the coherent subnetworks

There is a “core” of participating regions that recruits more regions by increasing the spectral coherence (or excludes regions by decreasing the coherence). As the coherence takes into account both zero-lag and constant-lag synchrony, the coherent functional subnetworks include distant regions that are phase-locked but not in-phase with each other (Prado et al., 2023b).

High activity of a region at a peak frequency, measured by the fractional occupancy of each region (Figure 2-figure Supplement 1), does not distinguish the presence of the region in multiple functional subnetworks. In fact, a region with a broader spectrum than a single peaky frequency, i.e., with a high spectral entropy, could be participating in more than one functional subnetwork instantaneously. For example, empirical data has demonstrated that sensory networks exhibit interaction of theta and gamma bands of EEG (***Canolty et al., 2006***; ***Samiee and Baillet, 2017***), relating to a bottom-up and a top-down functional network, respectively.

On the other hand, the fractional occupancy of each functional subnetwork (Figure 4) indicates the switching of the pattern of coherence for each frequency band. In this way, it offers a perspective of how the synchronization patterns distribute differently over time. We remark that as the fractional occupancy is a ratio over the total simulation period, the different time window length does not matter to qualitatively compare this metric result between frequency bands.

We found that for the two lower-frequency peaks there exists a difference in fractional occupancy between all subnetworks. Taking into account that the duration distribution of the subnetwork with higher fractional occupancy also has a longer tail, it indicates a temporal preference of the dynamics to remain in that subnetwork. Thus, at the lower frequency peaks the simulated brain signals show an “idle” mode, that as mentioned before, spatially resembles the DMN (***Callard and Margulies, 2014***). In contrast, there are no statistical differences for the occupancy at the two higher frequency peaks. Considering their resemblance with the executive control network, the lack of differences in fractional occupancy indicates that the resting brain does not prefer a cognitive task.

We use sliding windows containing several cycles for our coherence calculation. This meant that the median duration of each subnetwork is larger than the typical microstates duration of 50 ms (***Vidaurre et al., 2016***; ***Larzabal et al., 2023***). Reliable coherence measurements are possible with just a fraction of the cycle (***Vidaurre et al., 2018***), but we preferred to be conservative in the analysis. In any case, our subnetworks are concurrent at the different frequency peaks. The combined overlap of the subnetworks results in states with lower median duration than 50 ms (Figure 4-figure Supplement 1) but with an unfeasible number of possible clusters (*M*=3925). However, this is explained by the fact that microstates duration is based on transition models (Hidden Markov Model and variants (***Vidaurre et al., 2016***)), and the combined overlap of the coherent subnetworks reflects the dynamic of transitions between subnetworks.

## Conclusions

The increase of the dynamical richness by the human-connectome interaction of single intrinsic frequency oscillators is well described by the spectral entropy. We propose that spectral entropy is a proxy measurement of multi-state metastability, which is expected to be high at the awake resting-state brain. In line with this approach, the emerging coherent subnetworks at the point of maximal spectral entropy resemble the default mode network, the salience network, and the executive control network, not only in their spatial regions, but also near the respective well-known EEG brain rhythms. Further, the correlations fit with the average spectrum from other brain states EEG are located at points in the parameters-space of lower spectral entropy, suggesting a relation of this dynamical measurement of neural activity with behavior calm.

In addition, we found that the multiple coherent subnetworks at lower frequencies, resembling the DMN, have different fractional occupancy. On the other hand, the executive control-like subnetworks are similar in their fractional occupancy. However, the duration distributions have differences for almost all the frequencies. These findings indicate the presence of a default “idle” stage without preference to switch into a preferred executive subnetwork on the maximal spectral entropy dynamics. That was what we expected from the brain in resting-state, which makes the model suitable for its use as a baseline for the simulation of awake brain interventions.

## Methods and Materials

### Kuramoto Model

The Kuramoto model (KM) describes the synchrony dynamics of a harmonic oscillator coupled inside an oscillator network. The differential equation of KM describes the time evolution of the phase of one oscillator subjected to perturbations from neighboring oscillators (***Breakspear et al., 2010***). The simple Kuramoto model equation, with equal connection strengths and all-to-all coupling between oscillators, becomes:

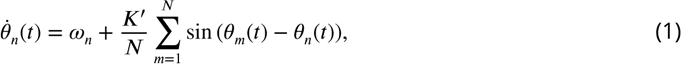

where *ω*_*n*_ is the natural frequency of the nth oscillator, *K*^′^ is the global coupling strength between all oscillators, *N* is the total number of nodes, and *θ*_*n*_ and *θ*_*m*_ are the phase shifts between two oscillators, respectively.

In order to use human connectome data, we employed a modified KM with heterogeneous coupling coefficients and delays between the connections. The model is represented by:

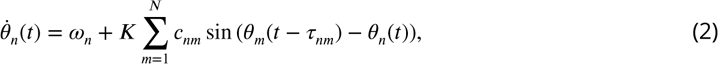

where *c*_*nm*_ is the element from the weighted adjacency matrix that represents the coupling strength, and *τ*_*nm*_ represents the delay. We simplified the term *K*^′^∕*N* from expression (1) to *K*, as *N* = 90 is the same for all simulations. We used the AAL90 (***Tzourio-Mazoyer et al., 2002***) connectivity matrices with volume correction, as in ***Cabral et al. (2011***, ***2014***).

As an additional global parameter, we also vary *τ*_*nm*_ by a mean delay factor MD. To achieve this modulation, the matrix *τ* of delays is normalized by the average of the nonzero elements and then multiplied by the mean delay factor. As our connectome data is a distance matrix between brain regions, the applied mean delay is a proxy for the inverse of the mean conduction speed (***Cabral et al., 2011***).

We simulate 300 seconds for each set of the parameters’ space in Figure 1. We perform twenty simulations, each of 30 minutes, with different initial conditions for *θ*_*n*_ using the point of maximal spectral entropy.

Structural features of human-connectome

AAL90 consists of two non-negative matrices: one matrix, *C*, of the connection weights with elements *c*_*nm*_, and another matrix, *D*, of the distances between regions.

The matrix *D* is symmetric, but the matrix *C* is not. Then, we calculate the connectivity strength of each region/node *n* as if we use a symmetric version of the *C* matrix by averaging the directed weights using *c*^′^ = (*c*_*nm*_ + *c*_*nm*_)∕2 (***Deco et al., 2017***), then

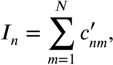

and the node degree as:

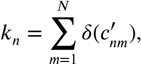

where the function τ(*c*^′^) is 1 if *c*^′^ >=1 > 0 and zero otherwise.

To vary the mean delay, the *D* matrix is normalized by its average taking into account only the values where the *c*_*nm*_ > 0. Then, *τ*_*nm*_ is defined by

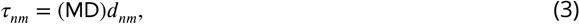

where *d*_*nm*_ is an element of the normalized *D* matrix.

### Dynamic measurements on simulated signals

#### Kuramoto order parameter

The measure of the global synchronization between oscillators as a function of time uses the average phase value, the Kuramoto order parameter (KOP) (***Breakspear et al., 2010***), expressed by:

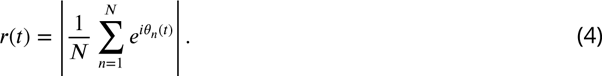

#### Frequency spectrum

All the nodes in the KM have an intrinsic frequency (*ω*_*n*_). The intrinsic frequency of auditory evoked responses of a vast amount of cortical regions is around 40 Hz (***Pastor et al., 2002***), and this intrinsic frequency was previously used in ***Cabral et al. (2014)***. Thus, we use *ω*_*n*_ = 2π(40) for all nodes. However, inter-node interactions may cause individual phase shifts leading to changes in the frequency of each node and the frequency of the system as a whole (***Cabral et al., 2014***).

Then, the spectrum of each oscillator could have broader bandwidth with additional frequency peaks. We calculate the power spectral density of each node using a windowed Fourier transform of sin (*θ*_*n*_(*t*)) via Welch’s method. We use a Hann time window of 5 seconds, overlapping by 50%, to get a frequency resolution of 0.2 Hz.

#### Spectral Entropy

The normalized power spectral density by the sum of the power along all the frequencies could be considered a probability distribution function. Also, taking advantage of the quantization of the frequency domain in discrete bins by the numerical calculation of the Fourier transform, we take the Shannon entropy of the normalized power spectral density (Inouye et al., 1991), given by:

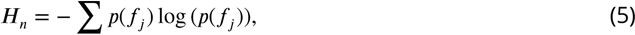

where *f*_*j*_ indicates the *j*-th frequency bin, and *p*(*f*_*j*_) the normalized power spectral density at that frequency bin. Further, we sum the spectral entropy over all the nodes.

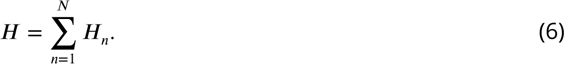

Selection of frequency peaks

The frequency peaks are the bins for which the average spectrum (taken over all nodes) has a local maximum. We selected five frequency peaks for the set of parameters {*K* = 4, mean delay = 21*ms*} that correspond to the set **p** = {13, 15, 29.5, 41.25, 42.75} Hz.

#### Selection of envelopes threshold

A proxy to the instantaneous power of each node at a particular frequency was calculated with the envelope of the narrow-band filtered signals. This comes from the amplitude or magnitude of the Hilbert transform of the filtered signals. To discretize the time a node occupies in each frequency peak, we employed a binary classification approach with a single threshold for all the 90 nodes and the 5 frequency-band envelopes. Then, we select the threshold of the maximum F1-score considering: *True Positives:* time samples where just one envelope overpass the threshold; *False Positives:* time samples where more than one envelope overpass the threshold; *Negatives:* time samples where none of the envelopes overpass the threshold.

### Functional connectivity matrices

The FC matrices were calculated with the spectral coherence between the nodes’ activity (***Lowet et al., 2016***). The time window length comes from expecting to include at least six complete oscillations. Then, the time windows were {462, 400, 204, 146, 140} ms for the correspondent peak frequency. The spectral densities calculation employs shorter windows of three complete oscillations with an overlap of 50%. In addition to disentangling the networks that co-occur but have different phase synchrony, we also calculated the FC matrices with the imaginary part of the coherence.

### The spectral coherence is defined by

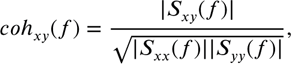

where *S*_*xy*_(*f*) is the cross-spectral density of signal *x* and signal *y*. *S*_*xx*_(*f*) and *S*_*yy*_(*f*) are their respective spectral densities.

The imaginary part of the coherence (***Nolte et al., 2004***; ***Hinkley et al., 2013***) is defined by

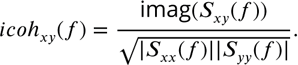

#### FC matrices clustering

The FC matrices are symmetric, and all diagonal elements are one. Then, before clustering, we re-arrange the lower triangular matrix excluding the diagonal as a vector. This gives an array *FC*^(*s*)^(*p*) ∈ *R*^*w*×4005^, where *W* is the number of time windows, for each trial *s*, and for each frequency peak in **p**.

We used the SpectralClustering method from the Sckitlearn Python library, with ’nearest_neighbors’ as the affinity parameter to cluster the data. This algorithm was performed over the PCA transformation of the FC array data using ten components. Before using the clustering algorithm, we normalized the PCA-transformed data by subtracting the minimum value and then dividing it by the difference between the maximum and the minimum value.

The number of clusters, *M*, is a parameter of the clustering algorithm. We employed two criteria to select the quantity of the clusters. The first is the inter-trial correlation between the most similar centroids of the clusters from each trial (Figure 3-figure Supplement 5 A).

The second is the correlation of the cluster size between trials, which corresponds with the fractional occupancy of each cluster (Figure 3-figure Supplement 5 B). This second criterion reduces the search range to *M* < 11.

In particular, we selected the number of clusters, *M*, that is the maximum of the inter-trial correlation for the peak frequencies 13 Hz, 15 Hz, and 29.4 Hz. For the peak frequencies 41.2 Hz and 43 Hz, we selected *M* = 7, just before the ’elbow’ at *M* = 8.

The subnetworks shown in Figure 3 correspond to the centroid of the clusters. For visualization, we pruned the not-present edges in the structural matrix *C*, the edges with a standard deviation intra-cluster above the 80% percentile, and the edges with coherence below the 90% percentile, this is keeping the 10% of connections with higher coherence.

## Acknowledgments

This work was supported by Agencia Nacional de Investigación y Desarrollo de Chile (ANID): FONDECYT Postdoc No. 3230682 (FAT), FONDECYT Postdoc No. 3210508 (MO), FONDECYT 1201822 (WED), FONDECYT 1231132 (AW), Anillo ACT210053, and Basal FB0008, BASAL FB210008 (MO); La Caixa Junior Leader Fellow (JC).

---

**Figure 1—figure supplement 1.**
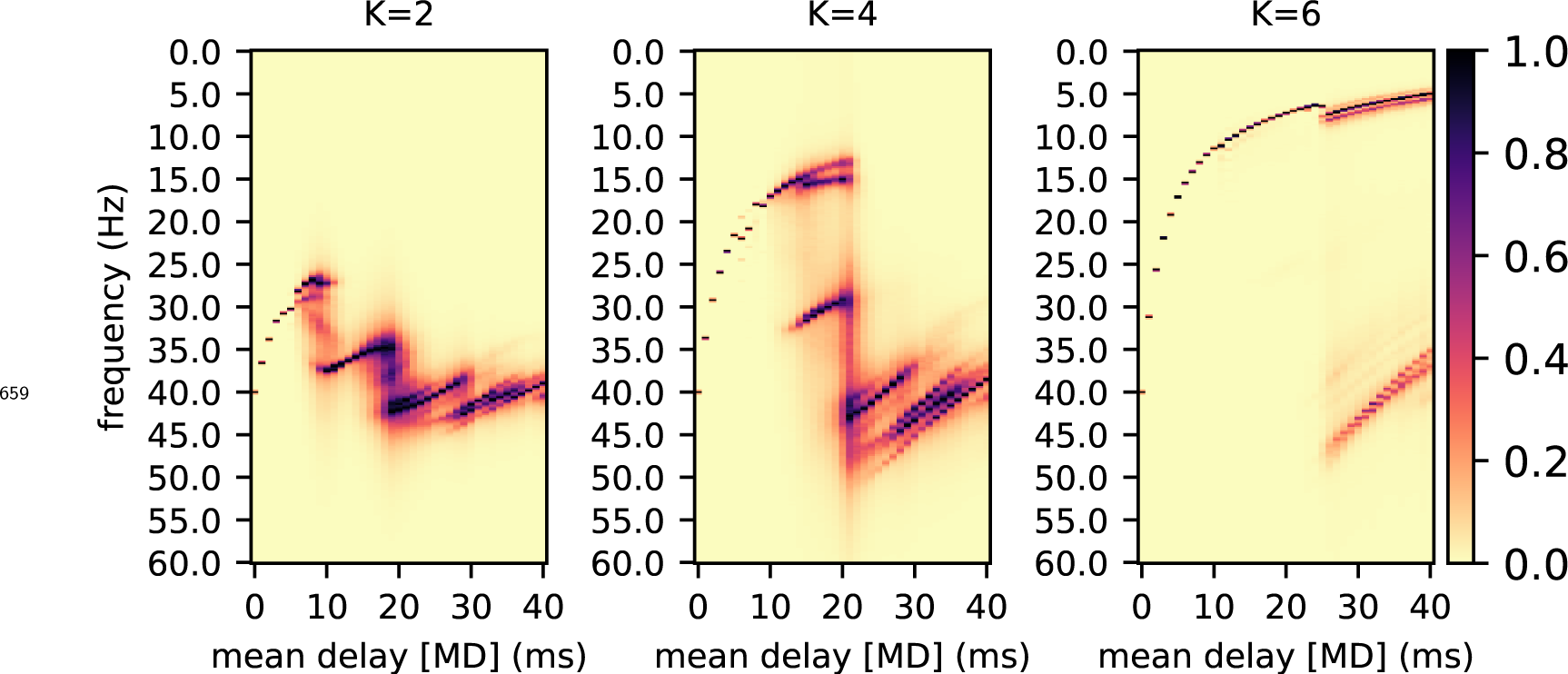
Normalized average spectrum for three values of *K* = {2, 4, 6} and for different values of mean delay.

**Figure 1—figure supplement 2.**
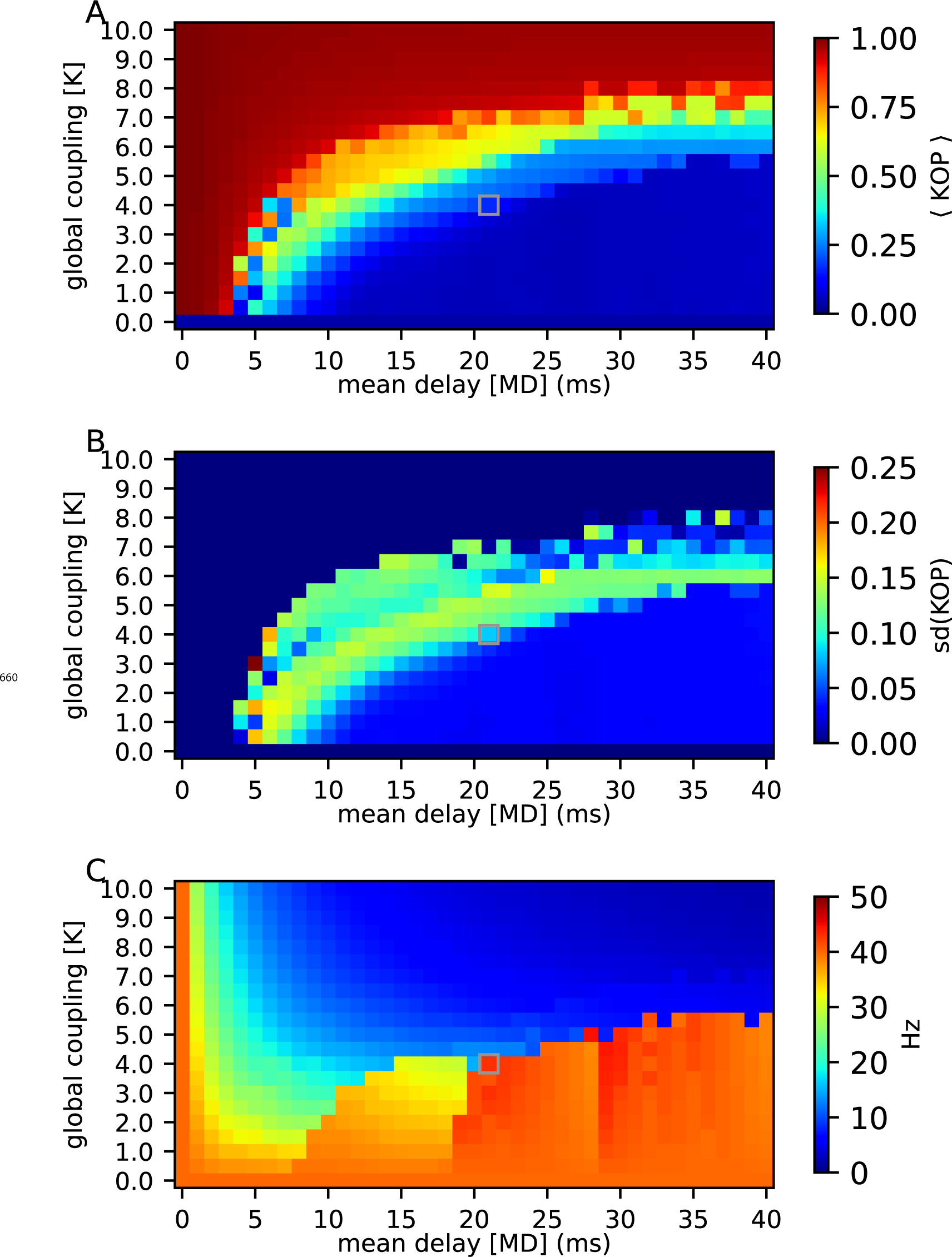
A. Average Kuramoto order parameter, the synchrony of the entire system. B. The standard deviation of the Kuramoto order parameter as another metastability metric. C. Peak frequency of the average spectrum.

**Figure 2—figure supplement 1.**
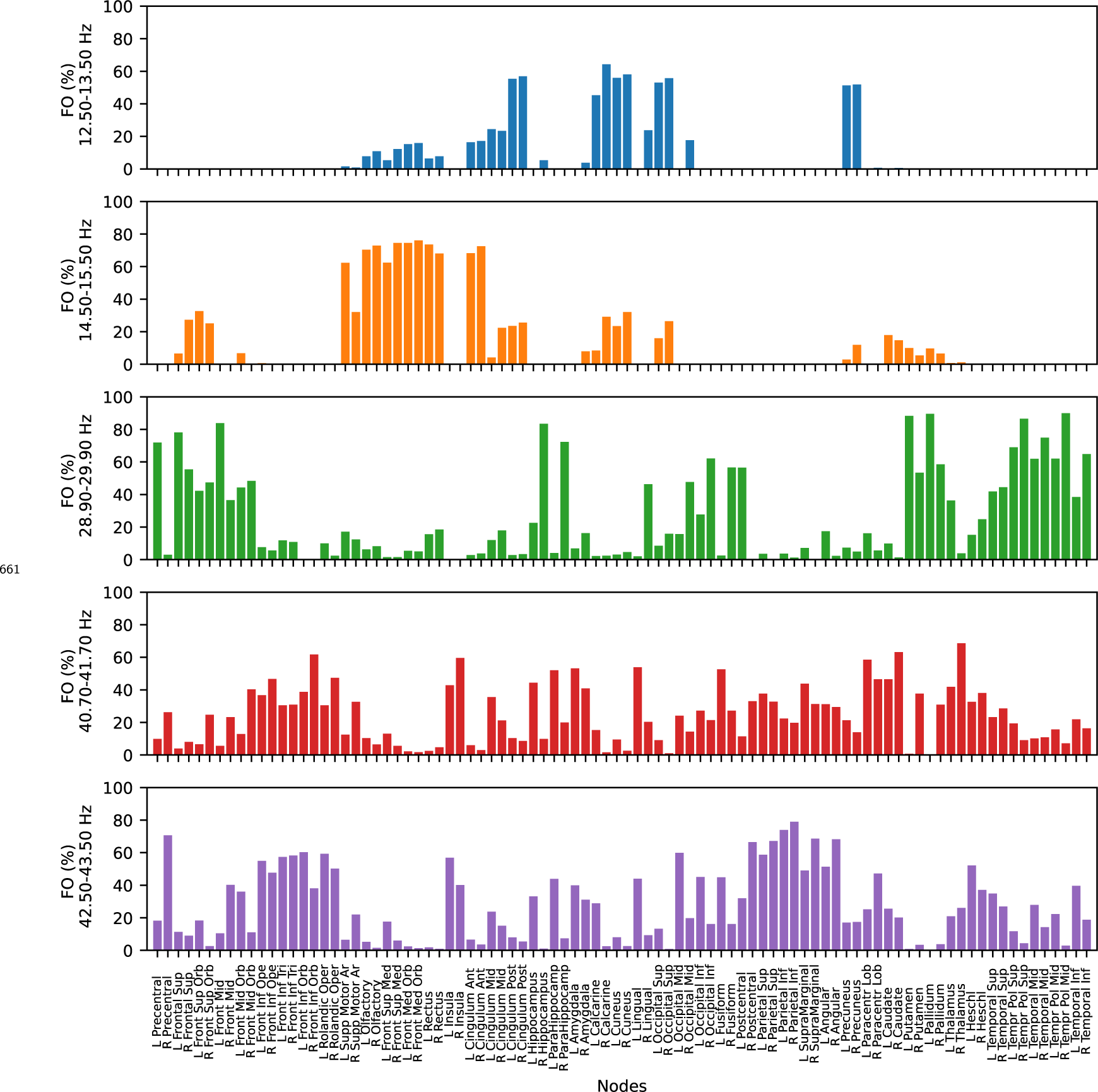
Fractional occupancy of the ninety nodes at each frequency band.

**Figure 2—figure supplement 2.**
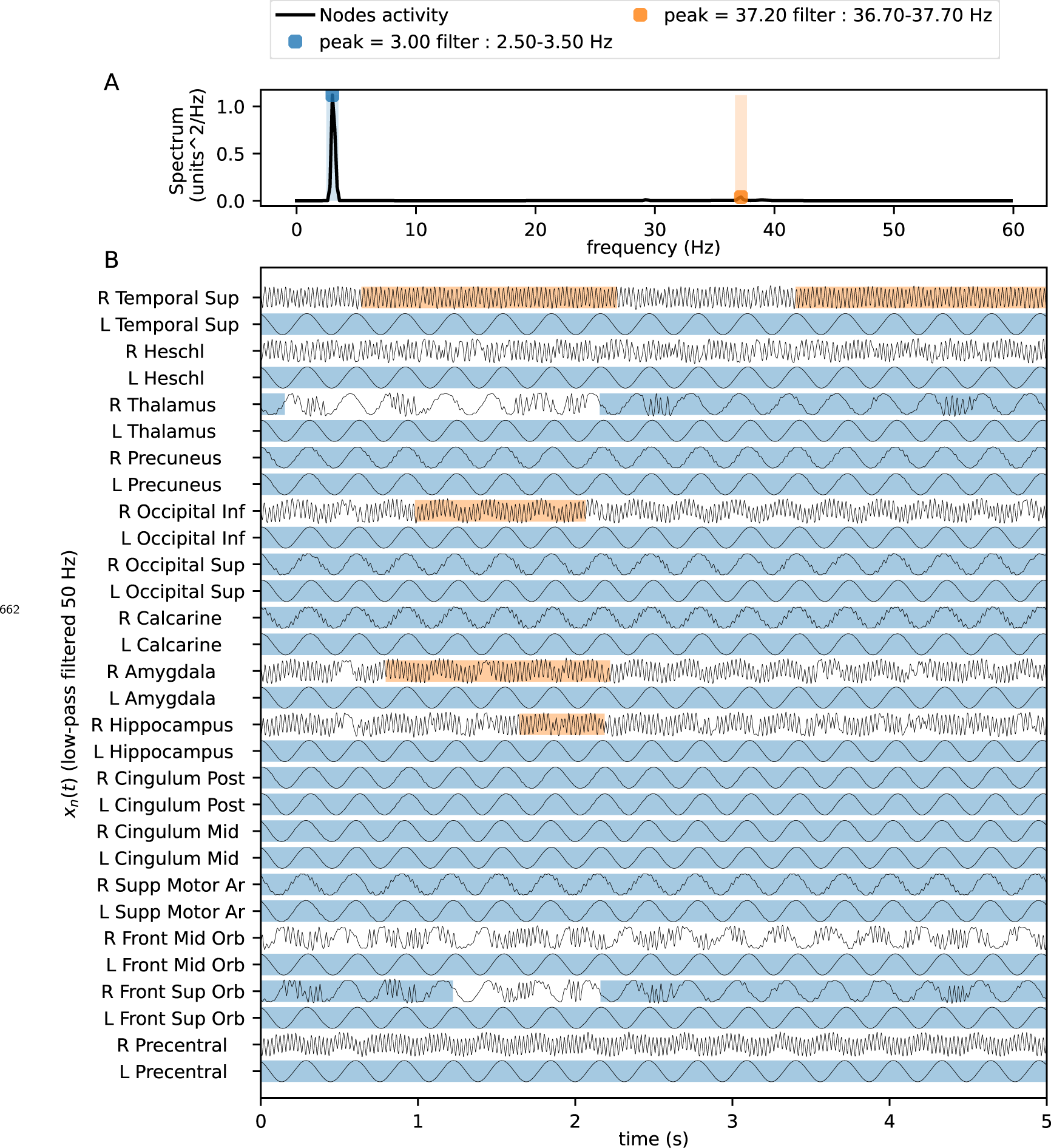
Average spectrum and time-series of the nodes using *K*=8, MD=37 ms.

**Figure 3—figure supplement 1.**
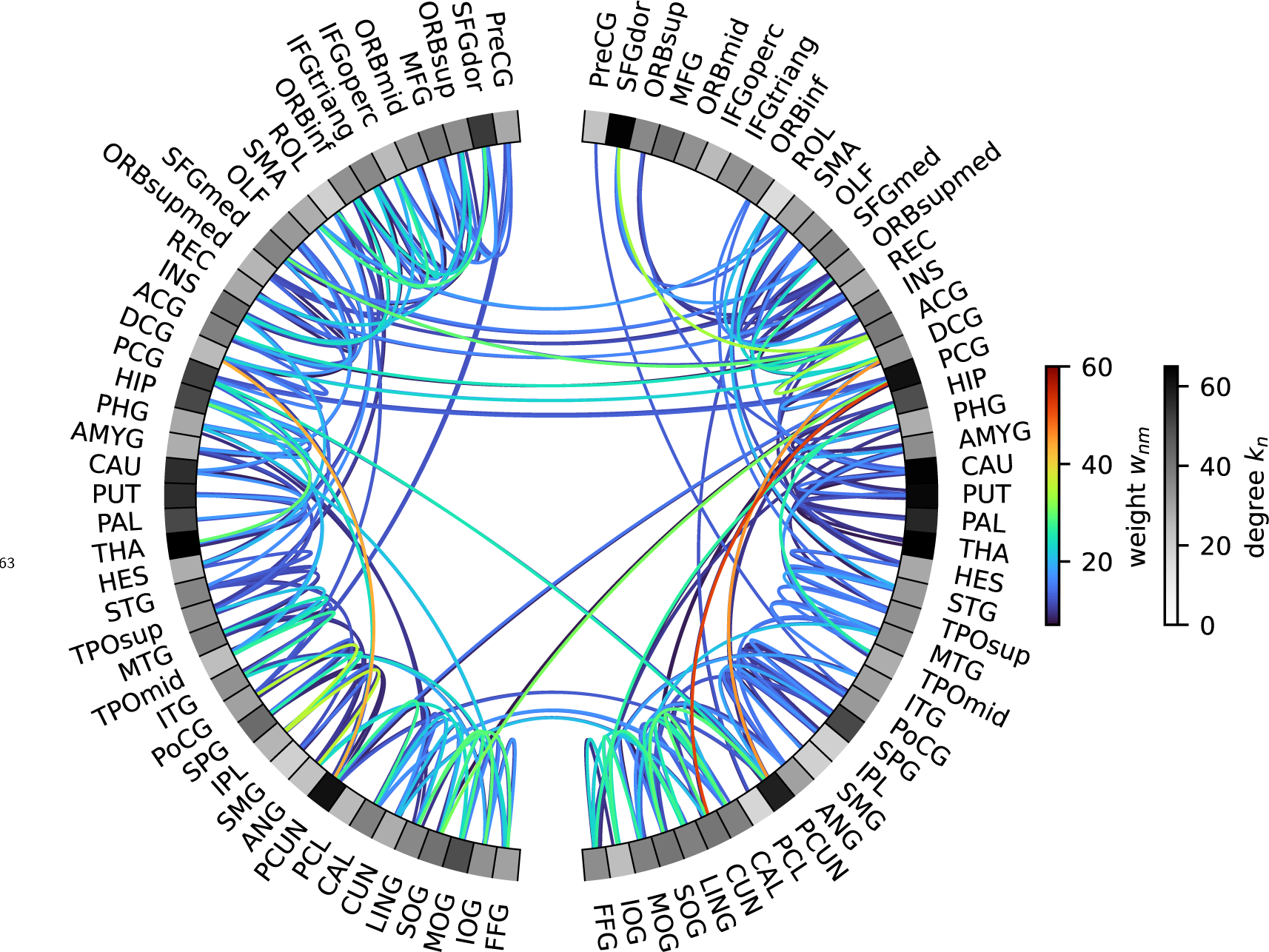
Structural connectivity and region degrees. The region labels correspond to the region in the same position of *Figure 3* and *Figure 3*-*figure Supplement 2*. The plotted edges are only the 10% higher connection weights. The gray color level of each region corresponds to the region degree.

**Figure 3—figure supplement 2.**
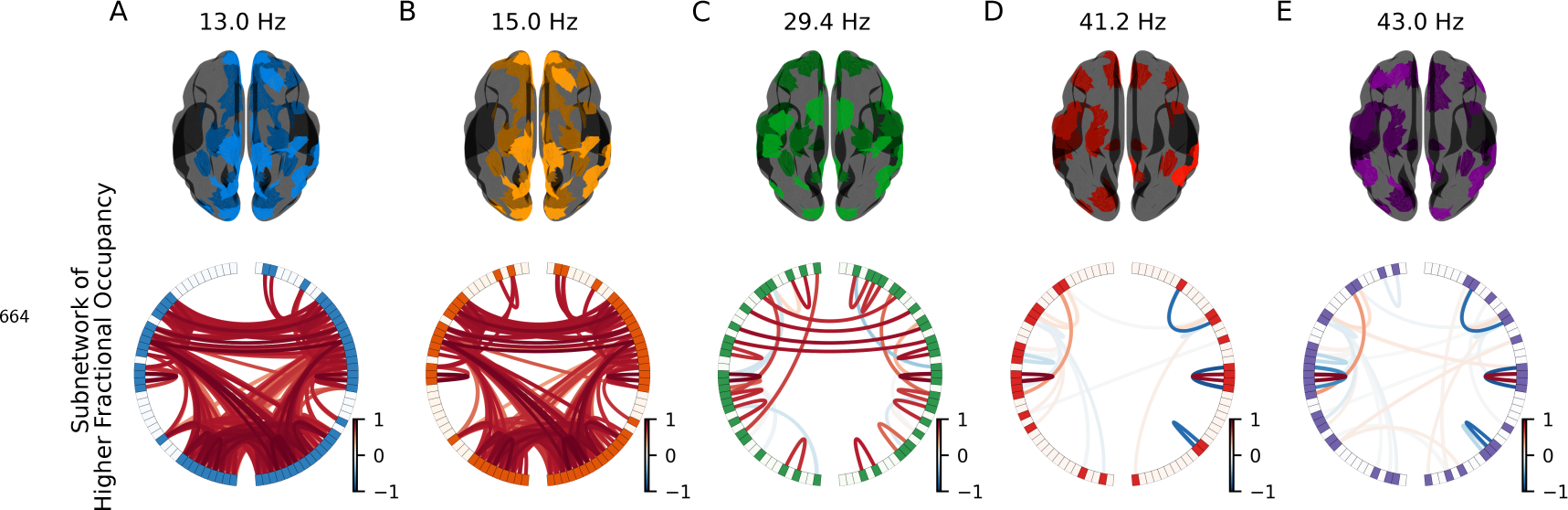
Real part of the edges coherence. The edges of the functional subnetworks have lower coherence as the frequency increases.

**Figure 3—figure supplement 3.**
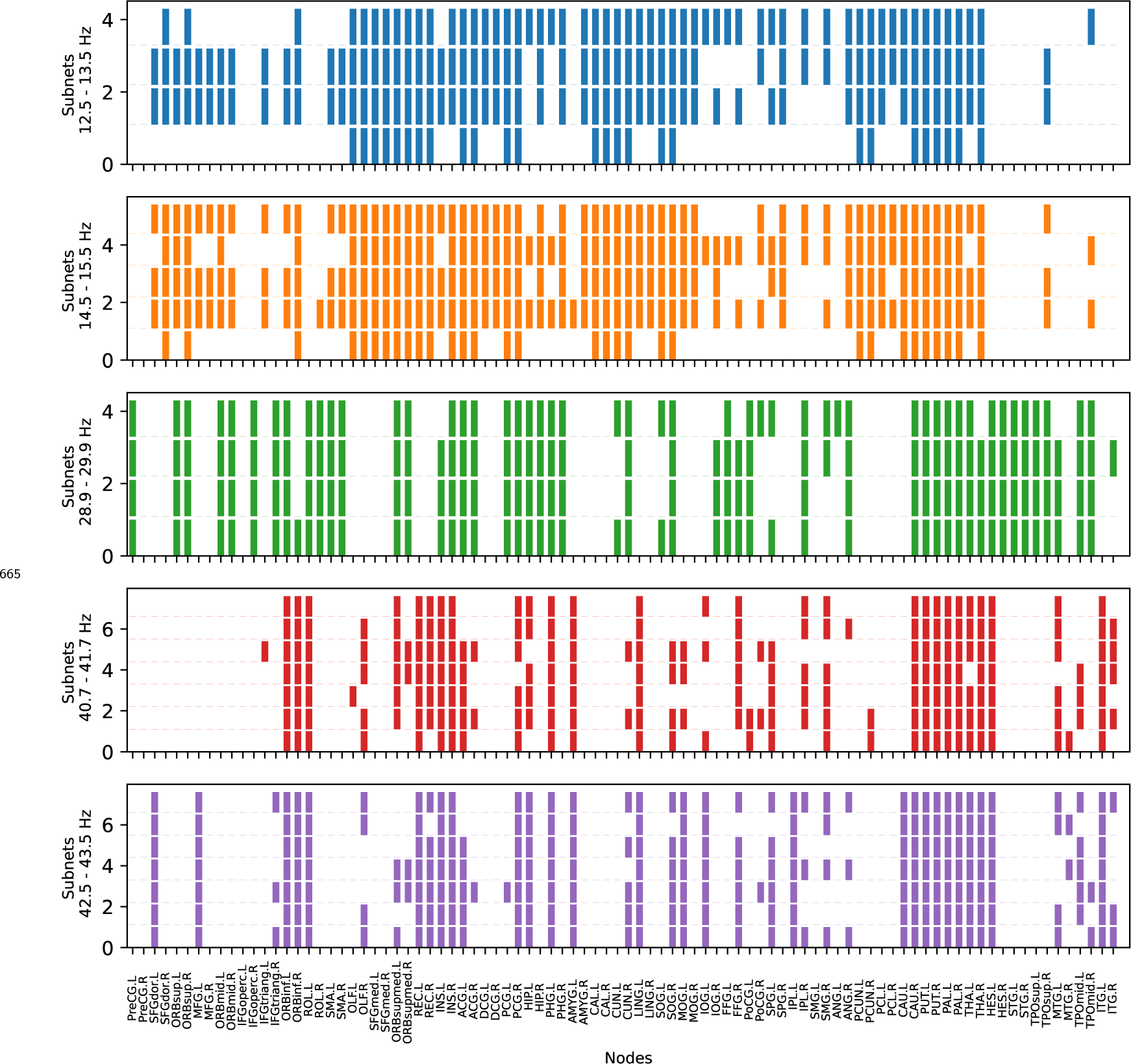
Regions participating in all the obtained subnetworks. Using the original node order of the AAL90 matrix.

**Figure 3—figure supplement 4.**
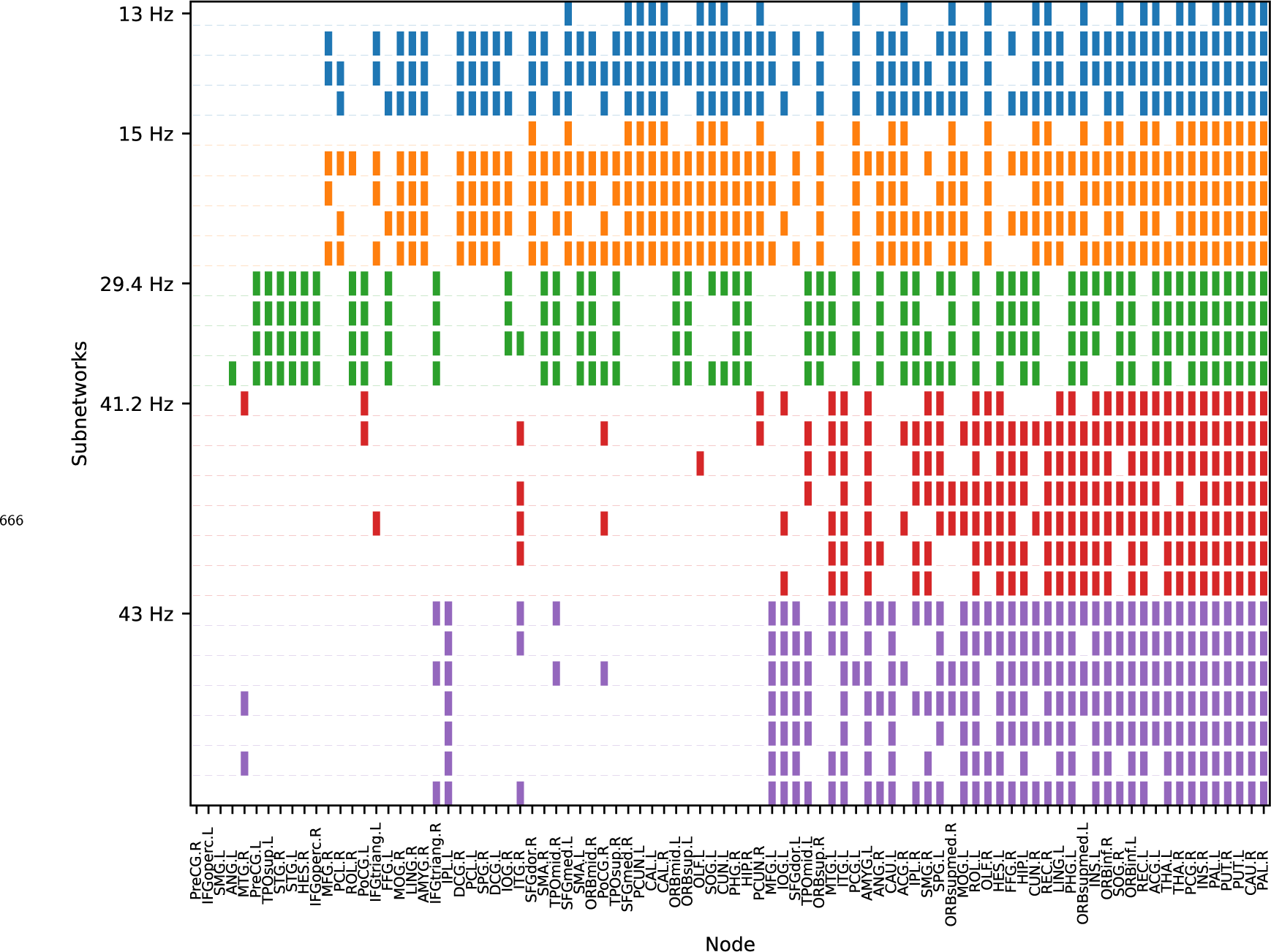
Regions participating in all the obtained subnetworks, sorted from the least to the most functional connected node.

**Figure 3—figure supplement 5.**
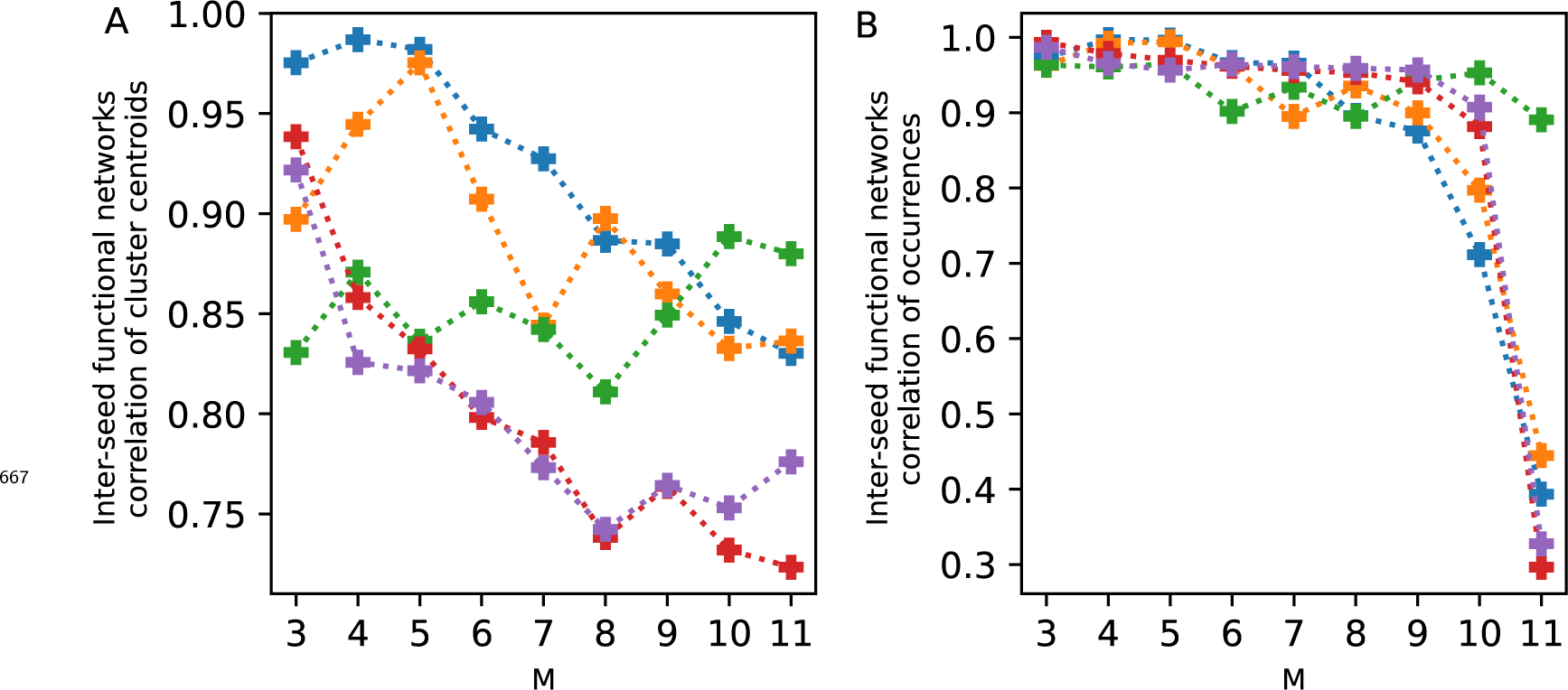
Clustering score. The two clustering criteria are detailed in Methods and Materials section. The number of clusters for each frequency peak is the.. value with the highest inter-trial correlation of the clusters centroids or where an elbow occurs. We also keep a high inter-trial correlation of the fractional occupancy. We selected 4, 5, 4, 7, and 7 subnetworks for the 13, 15, 29.4, 41.2, and 43 Hz frequency peaks, respectively.

**Figure 4—figure supplement 1.**
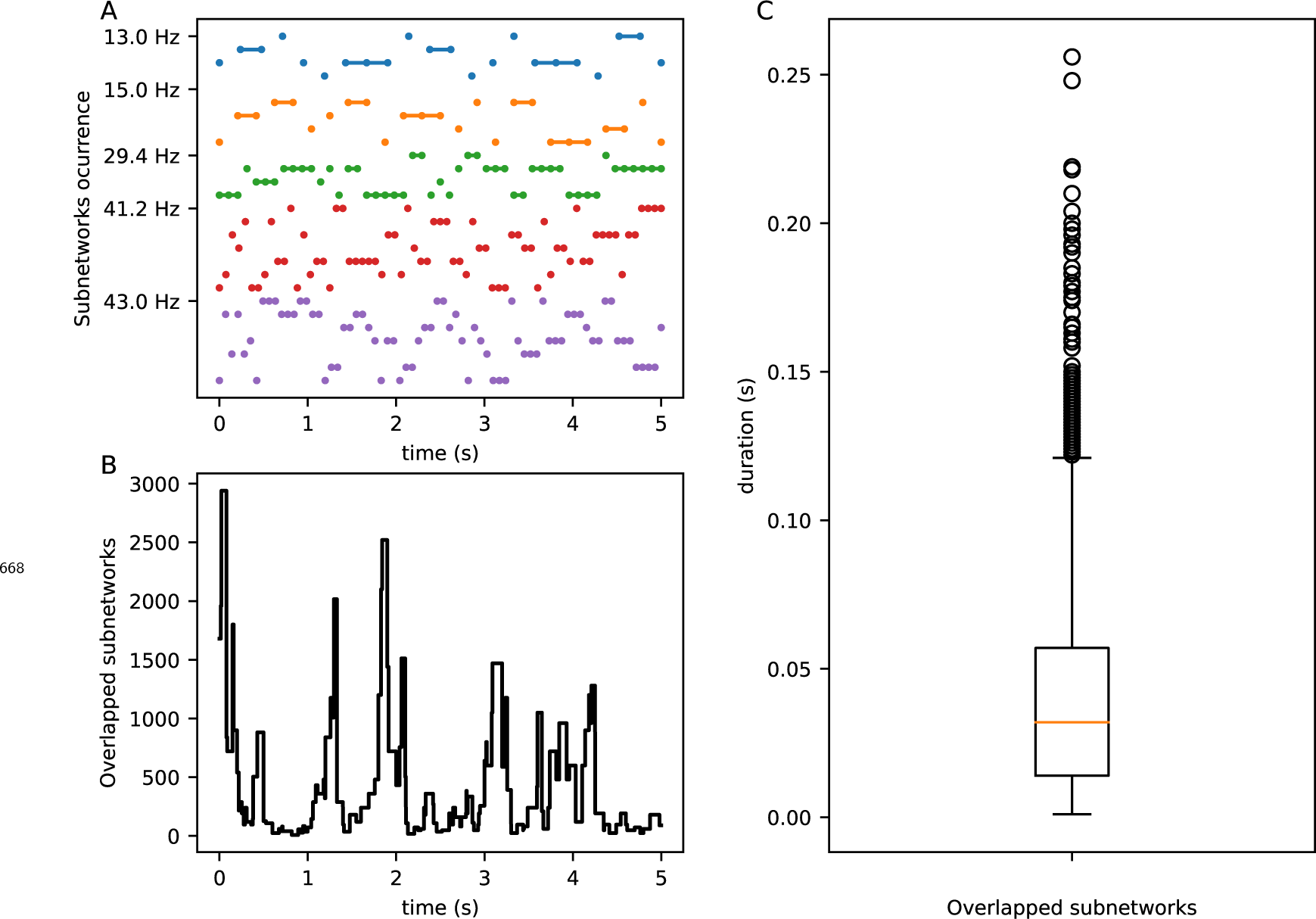
Excerpt of subnetworks occurrence. A. Occurrence of the subnetworks at each peak frequency along time. At each time point, there is one subnetwork of each peak frequency. B. Single time series of the overlapped subnetworks by relabeling the co-occurrences (There are a possible 3920 combinations from the 26 subnetworks at the five peak frequencies). C. Distribution of durations for overlapped subnetworks.

